# Intracellular Sodium Regulates Opioid Signalling in Peripheral Neurons

**DOI:** 10.1101/117952

**Authors:** Alexandros H. Kanellopoulos, Jing Zhao, Edward C. Emery, John N Wood

**Author notes:** Summary; intracellular sodium regulates neuronal GPCR signalling.

## Abstract

Opioid receptors signal more effectively in sensory neurons from pain-free mice lacking the voltagegated sodium channel Na_v_1.7. Type-A GPCRs are known to be regulated through a specific sodium binding site, the occupancy of which diminishes agonist binding. We have used an electrophysiological assay of Protein Kinase A activity to examine the role of intracellular sodium on opioid signalling. Phosphorylation of sodium channel Na_v_1.8 by activation of Protein Kinase A with db-cAMP is unaffected by altered intracellular sodium. By contrast, there is a dose-dependent inhibition of fentanyl action on Na_v_1.8 currents when intracellular sodium is increased from 0 mM to 20 mM. Fentanyl shows a 50% loss of activity and 80-fold increase in EC_50_ with 20 mM intracellular sodium. These data demonstrate that altered intracellular sodium levels modulate opioid receptor signalling.

---

Pert and Snyder showed the influence of sodium on opioid receptor activity in 1974, demonstrating that increased sodium concentrations caused diminished agonist binding (*1*). Forty years later the binding site for sodium on the δ-opioid receptor was identified by Fenalti et al. (*2*). Opioid receptors are members of the Class-A GPCR family that comprises about 700 members. A sodium binding site has been identified in adenosine A2_A_, β-1 adrenergic and PAR-1 receptors using ultra high resolution crystallography; a related site is present in almost all Class-A GPCRs (3-5). Modelling studies of the A2_A_ receptor have suggested that sodium may access receptors via a water-filled passage that links extracellular and intracellular sides of the receptor (*6*). Extracellular sodium is maintained at about 145 mM, whilst intracellular sodium is very much lower, in the range of 5 mM for sympathetic neurons (*7*) to 15 mM in lobster nerves (*8*). What then is the physiological role of allosteric sodium binding when such binding sites should, in principal, be fully occupied? We have examined the effect of changing intracellular sodium concentrations on the activity of μ-opioid receptors in individual mouse sensory neurons using a sensitive electrophysiological assay. Protein Kinase A (PKA) is known to phosphorylate five serine residues in the first intracellular loop of Na_v_l.8, a tetrodotoxin-insensitive voltage-gated sodium channel that is uniquely expressed in sensory neurons (*9*). This results in a large increase in TTX-resistant sodium channel activity that can be quantitated by electrophysiological recording. Fentanyl, acting through μ-opioid receptors and Gi proteins can suppress the activity of PKA, and diminish the level of TTX-resistant (TTX-r) sodium current (9*,10,11*). These observations provide us with a simple assay system for measuring opioid action in intact cells, allowing us to vary the level of intracellular sodium and examine the consequences (Figure 1).

**Figure 1.**
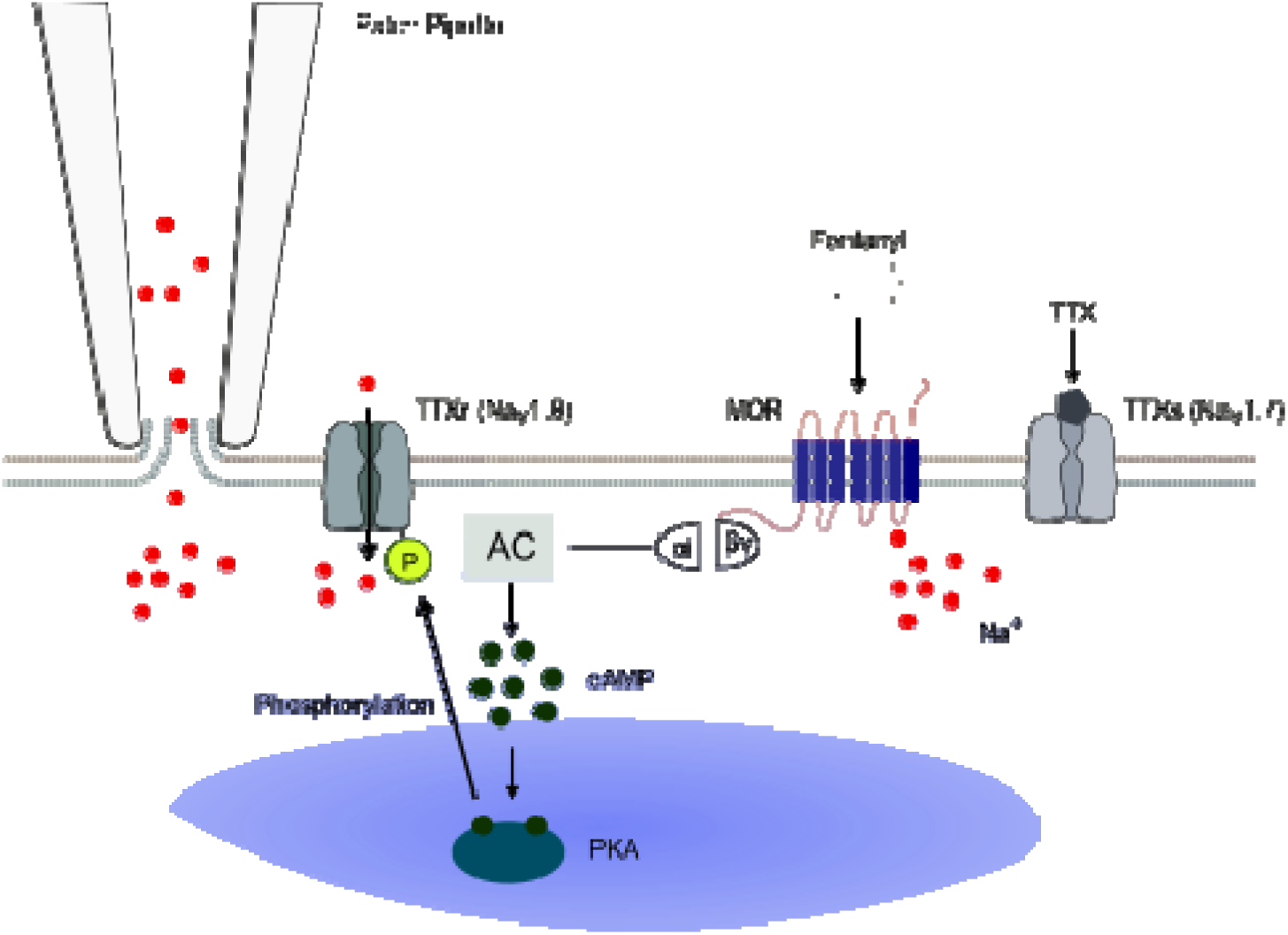
TTX-resistant currents were recorded using whole cell patch clamp recordings in dorsal root ganglion neurons, permitting the manipulation of intracellular sodium concentrations. TTX-sensitive currents were blocked using TTX. Opioid receptor-mediated Gi activity causes an inhibition of adenylyl cyclase leading to deceased PKA activity and lower levels of Na_v_l.8 phosphorylation, thereby decreasing sodium currents entering through Na_v_l.8. An accurate recording of opioid receptor activity following the exposure to fentanyl is acquired through measurement of TTX-resistant currents.

Intracellular sodium levels can be effectively altered in intact cells by perfusion from a patch-clamp recording electrode (*12*). First, we investigated the effects of fentanyl on Na_v_1.8-mediatedTTX-r sodium currents in sensory neurons (11). We found that there was a concentration-dependent inhibition of fentanyl-evoked inhibition of TTX-r currents with increasing intracellular sodium. Within the physiological range, 20 mM sodium substantially lowered the inhibitory activity of fentanyl on TTXr current density, whilst nominal 0 mM sodium concentrations potentiated inhibitory activity (Figure 2).

**Figure 2.**
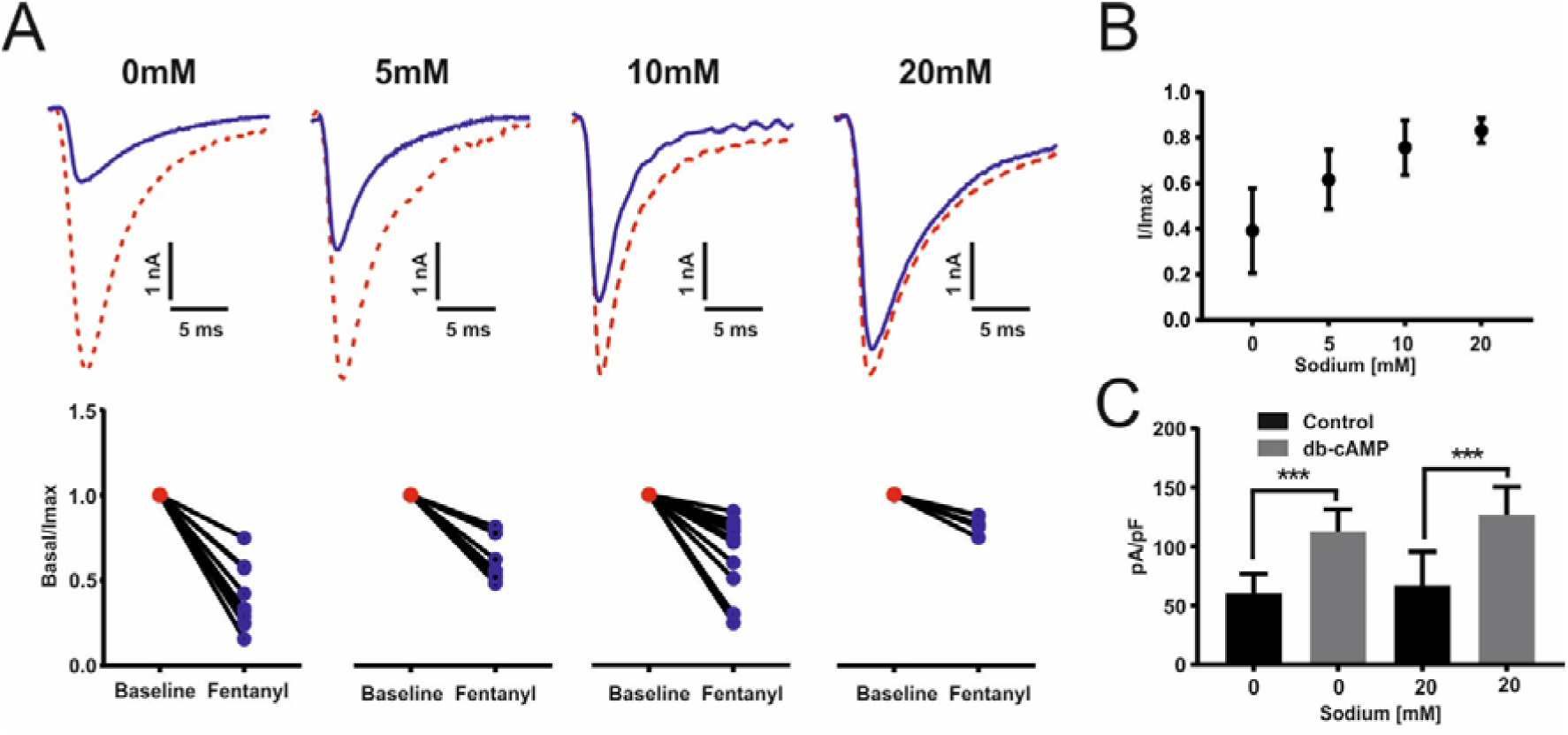
(A) Electrophysiological example traces of TTX-resistant Na_v_1.8 currents from dorsal root ganglia neurons following the exposure to 100 nM fentanyl at different intracellular concentrations of sodium (0 mM, 5 mM, 10 mM and 20 mM). Corresponding dot plot of TTX-resistant Na_v_1.8 currents following the exposure to 100 nM fentanyl at different intracellular concentrations of sodium (0 mM: n=14, 5 mM: n=10, 10 mM: n=14 and 20 mM: n=6. All currents are normalised and compared to baseline. Student’s *t* test, 0 mM p<0.001, 5 mM p<0.001, 10 mM p<0.001 and 20 mM p<0.05. (B) Data from (A) plotted as normalised peak currents compared to sodium concentration. Data represents ±SEM. (C) Electrophysiological recordings of TTX-resistant Na_v_1.8 currents in dorsal root ganglia neurons following exposure to db-cAMP in 0 mM and 20 mM intracellular concentrations of sodium. WT 0 mM vs db-cAMP 0 mM, ***p<0.0002, change from baseline ∆=52.3 pA/pF. WT 20 mM vs db-cAMP 20 mM, ***p<0.0006, change from baseline ∆=59.8 pA/pF. No significant change between db-cAMP 0 mM vs db-cAMP 20 mM.

We next tested whether the downstream activity of PKA on Na_v_l.8 was modified by altered sodium levels. Using the PKA activator db-cAMP, we found that the level of TTX resistant Na_v_l.8 channel activity was upregulated by 53.4% in the presence of a nominal 0 mM intracellular sodium and by 52.8% in the presence of 20 mM sodium. There was no difference in db-cAMP-mediated effects on TTX-r currents in 0 mM or 20 mM sodium. Thus downstream sensitisation of Na_v_1.8 by PKA is independent of intracellular sodium concentration.

We then compared the activity of fentanyl on neurons expressing Na_v_1.8 with either 0 mM or 20 mM intracellular sodium. The maximal effect of fentanyl on Na_v_l.8 activity was lowered by 50% in the presence of 20 mM intracellular sodium. In addition, the EC_50_ for fentanyl action changed from 2 nM in 0 mM sodium to 126 nM in the presence of 20 mM sodium (Figure 3). Strikingly, these data recapitulate early studies of sodium effects on opioid binding carried out by Kosterlitz on brain membranes where a 40-fold loss of fentanyl activity was noted at high sodium conditions (13) compared to a 60 fold change in the functional assays described here. The simplest explanation of these data is that the activity of the μ-opioid receptor is regulated by intracellular sodium.

**Figure 3.**
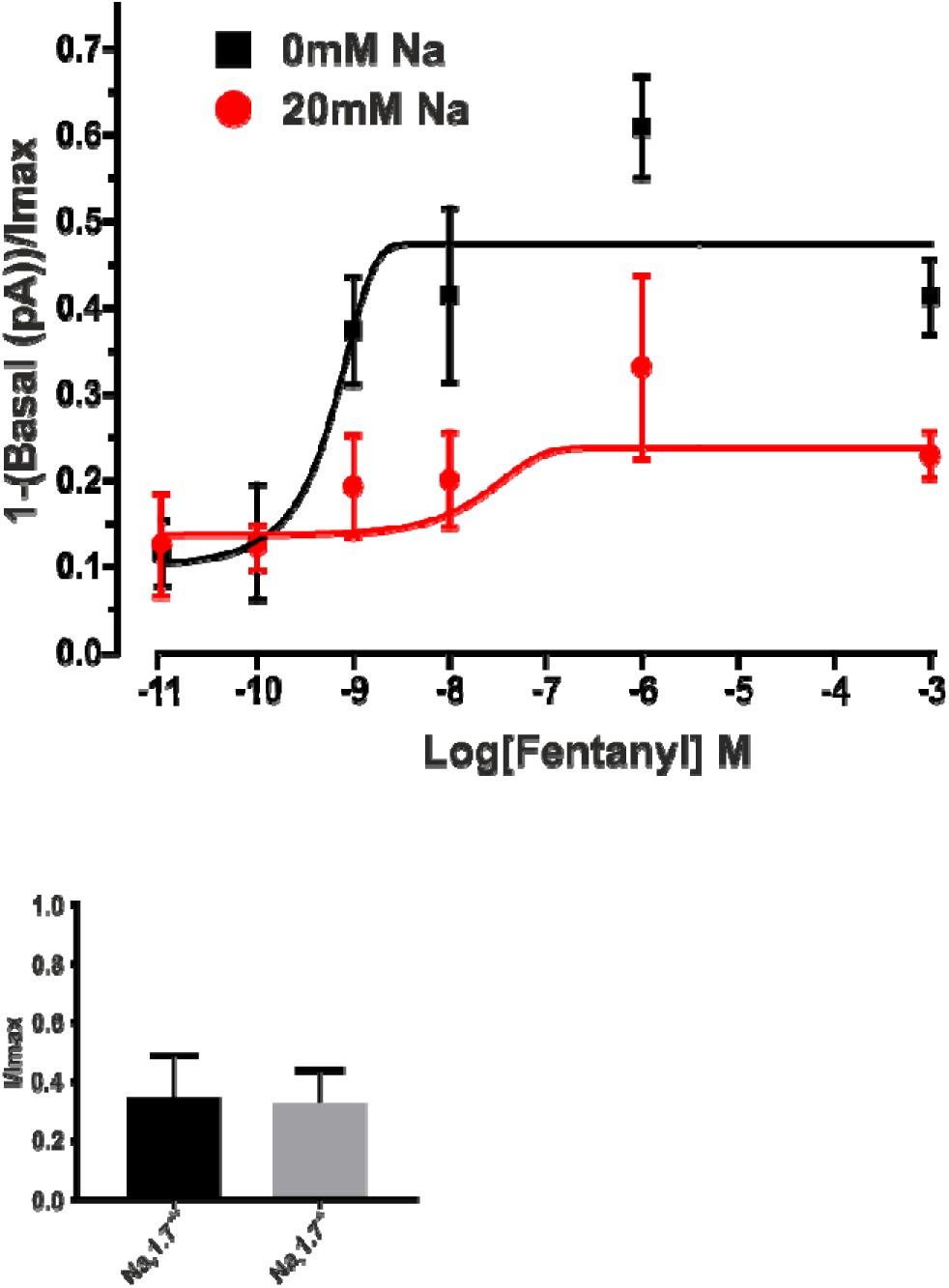
Fentanyl dose response curve. Electrophysiological recording following exposure to 1 pM, 10 pM, 100 pM, 1 nM, 100 nM, 1 mM concentrations of fentanyl at 0 mM and 20 mM intracellular sodium concentrations. All points for both 0 mM and 20 mM n=5. 0 mM EC_50_=2.12 nM, 20 mM EC_50_=0.129 μM. Maximal inhibition of PKA (65%) by fentanyl with 0 mM intracellular sodium occurs in both wild type and Nav1.7 null mutant sensory neurons. Responses are normalised to controls without fentanyl.

Does this event have biological significance? In earlier experiments, we found that the activity of PKA on Na_v_1.8 currents was dramatically diminished by fentanyl in Na_v_l.7 null mutant mice compared to wild type controls when intracellular sodium was held at 10 mM (11). These mice are pain-free, due in part to increased opioid drive (14). In addition, the effectiveness of fentanyl in blocking PKA is dramatically enhanced in Na_v_1.7 null mutant neurons using an immunohistochemical single cell assay of PKA phosphorylation (11). This would suggest that sodium entering through Na_v_1.7 ion channels regulates opioid signalling mechanisms. We tested the effect of fentanyl on wild type and Na_v_1.7 null mutant mice using different concentrations of intracellular sodium. At 0 mM sodium, the two strains showed identical maximal responses to fentanyl (Figure 3). However, at 10mM sodium, the Na_v_1.7 null mutant showed a more potent response than the wild-type control (11). This sodium concentration dependent effects is consistent with a potential role for sodium ingress through Na_v_1.7 as a second messenger in μ-opioid receptor regulation. How might this occur?

When epitope-tagged Na_v_1.7 is immunoprecipitated with channel-associated proteins from mouse DRG neurons, proteins that are known to bind to μ-opioid receptors also associate with Na_v_1.7, as judged by mass spectrometry. Thus GRIN1 that is associated with μ-opioid receptors also binds to Na_v_l.7 (Figure 4), and this interaction has been confirmed in a Western blot analysis of Na_v_l.7 immunoprecipitates (Fig 4). These findings provide a potential explanation for a specific effect of Na_v_1.7 activity on μ-opioid receptor function. By contrast, studies of Na_v_1.8 KO mice have shown that there is no effect on μ-opioid receptor activity as measured with a PKA assay following the deletion of this sodium channel gene (11). The physical apposition of Na_v_1.7 and μ-opioid receptors could explain the specific significance of sodium flux through Na_v_1.7 for the regulation of fentanyl signalling. Persistent currents associated with Na_v_1.7 have also been reported and these may have effects on local sodium concentrations near the ion channel (15).

**Figure 4.**
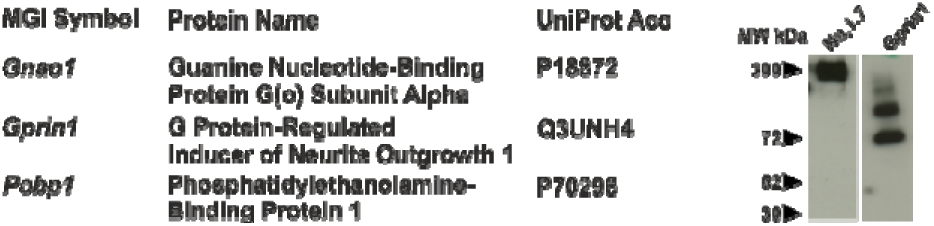
Table of identified proteins known to associate with both Na_v_1.7 and opioid receptors as identified using mass spectrometry. Western blots show that Na_v_l.7 co-immunoprecipitates with known opioid interactor Gprin1.

Does this observation have more general significance? If sodium is acting as a second messenger linking Na_v_l.7 activity with opioid GPCR signalling, it is possible that other cation channels could be linked to other class A GPCR receptor regulation in an analogous way. For example, adrenergic receptors are known to also regulate Na_v_1.8 through PKA mediated phosphorylation (16). In the case of Na_v_1.7, one may speculate that intense stimulation associated with extreme tissue damage could lead to the attenuation of opioid signalling and an amplification of nociceptor activity that could transiently potentiate avoidance behaviour. Whatever the biological consequences, the study presented here is consistent with a role for intracellular sodium as a second messenger controlling GPCR signalling.

## Acknowledgments

We thank the Wellcome Trust for their invaluable and generous support. AHK is a grand challenge UCL PhD student. AHK carried out the electrophysiological experiments, JZ and AHK carried out the interacting protein experiments, JNW and ECE conceived the study and JNW wrote the paper with input from all authors.

## Materials and Methods

### Neuronal Cultures

All animal experiments were approved by the United Kingdom Home Office Animals Scientific Procedures Act 1986. Experiments were conducted using both female and male mice. For experiments using transgenic mice, wild-type littermate animals were used as controls. All strains of mice used for procedures were of C57Bl/6 background. All mice used for experimentation were at least 6 weeks old. Sensory neurons were isolated as described in (14).

### Co-immunoprecipitation

Epitope-tagged Na_v_1.7 mice were generated, characterised and used for immunoprécipitation and mass spectrometric studies as described in Kannelopoulos et al. 2017. Nav1.7 complexes were purified with M2 Magnetic FLAG coupled beads (Sigma-Aldrich).

### Western blot

Samples for western blot were prepared by adding 3X Loading Buffer (Life Techniologies) containing DTT and denatured for 5 minutes at 95°C 30μg of samples were loaded in to precast gels (Bio-Rad) along with a multicolour spectra high range protein ladder (Thermo Scientific). Gels were placed in a gel tank and submerged in 1X Running Buffer. Samples were run for between 1-3 hours, depending on size of protein of interest at 120V. Immobilin-P membrane (Millipore) was activated with methanol (Sigma) following which wet transfer of proteins was done in 1X ice cold Transfer Buffer. Transfer was done for 1 hour at 100V. Membrane were then blocked using 5% Marvel in 1X PBS. Antibody incubation was then done overnight at 40C. The membrane was then washed and incubated with HRP tagged secondary antibody in PBS containing 25% skimmed milk, with agitation at room temperature. Membranes were then washed and the proteins were visualised using Super Signal West Dura Extended Duration Substrate (Thermo Scientific) on light sensitive film (GE Healthcare) and developed on a Konica Minolta (SRX-101A) medical film developer. Antibodies used were anti-HAT antibody LSBio LS-C51508 for Nav1.7tap-tag and GRIN1 Proteintech 13771-1-AP at 1;1000 dilution.

### Electrophysiology

All electrophysiological recordings were performed using an Axopatch 200B amplifier and a Digidata 1440A digitizer (Axon Instruments), controlled by Clampex software (version 10, Molecular Devices). Filamented borosilicate microelectrodes (GC150TF-7.5, Harvard Apparatus) were coated with beeswax and fire-polished using a microforge (Narishige) to give resistances of 2 to 3 megohms. For voltage-clamp experiments, the following solutions were used. The extracellular solution contains 70 mM NaCl, 70 mM choline chloride, 3 mM KCl, 1 mM MgCl2, 1 mM CaCl2, 20 mM tetraethylammonium chloride, 0.1 mM CdCl2, 300 nM TTX, 10 mM Hepes, and 10 mM glucose (pH 7.3) with NaOH. The intracellular solution contains 140 mM CsF, 1 mM EGTA, 10 mM NaCl, and 10 mM Hepes (pH 7.3) with CsOH. Unless otherwise stated, standard whole-cell currents were acquired at 25 kHz and filtered at 10 kHz (low-pass Bessel filter). After achieving whole-cell configuration, the cell was left for 5 min to dialyze the intracellular solution. A holding potential of −100 mV was applied, and series resistance was compensated by ≥70%. All currents were leak-subtracted using a p/4 protocol. To record TTXr sodium currents, we applied a depolarizing voltage-pulse protocol to the cell; the cell was held at −100 mV and then stepped to −15 mV for 50 ms before returning back to −100 mV. This step was applied every 5 s for the duration of the experiment. The cells were continuously perfused using a gravity-fed perfusion system. All electrophysiological data were extracted using Clampfit (version 10, Molecular Devices) and analyzed using GraphPad Prism software (version 6, GraphPad).

### Statistical Analysis

Statistical analysis were performed with either Student’s *t* test with respective post hoc tests. *P* < 0.05 was considered statistically significant. Voltage-clamp experiments were analysed using cCLAMP software and Origin (OriginLab Corp., Northampton, MA) software programs. Current density-voltage (pA/pF) analysis by measuring peak currents at different applied voltage steps and normalised to cell capacitance. Voltage dependent activation data was fitted to a Boltzman equation y = (A2 + (A1 − A2)/(1 + exp((Vh − x)/k))) * (x − Vrev), where A1 is the maximal amplitude, Vh is the potential of half-maximal activation, x is the clamped membrane potential, Vrev is the reversal potential, and k is a constant. All Boltzmann equations were fitted using ORIGIN software. ±SEM data were assumed to be normally distributed. Unpaired Student’s t test was used for statistical comparisons. Significance was determined at p < 0.05. Individual p values are given for each comparison made. Fentanyl dose response curve and IC50 calculations were fitted and measured using ORIGIN software.

